# *Verticillium dahliae* LysM effectors differentially contribute to virulence on plant hosts

**DOI:** 10.1101/085894

**Authors:** Anja Kombrink, Hanna Rovenich, Xiaoqian Shi-Kunne, Eduardo Rojas-Padilla, Grardy C.M. van den Berg, Emmanouil Domazakis, Ronnie de Jonge, Dirk-Jan Valkenburg, Andrea Sánchez-Vallet, Michael F. Seidl, Bart P.H.J. Thomma

**Affiliations:** Laboratory of Phytopathology, Wageningen University, Droevendaalsesteeg 1, 6708 PB Wageningen, The Netherlands.

**Author notes:** These authors contributed equally (Hanna Rovenich, Xiaoqian Shi-Kunne, Eduardo Rojas-Padilla, Grardy C.M. van den Berg). These authors contributed equally (Michael F. Seidl, Bart P.H.J. Thomma). Author for correspondence: Bart P.H.J. Thomma, Laboratory of Phytopathology, Wageningen University, Droevendaalsesteeg 1, 6708 PB Wageningen, The Netherlands.

**Keywords:** Vascular wilt disease, aggressiveness, tomato, Arabidopsis, tobacco, comparative genomics, chitin, hyphal protection, lineage-specific effector

## Abstract

Chitin-binding LysM effectors contribute to virulence of various plant pathogenic fungi that are causal agents of foliar diseases. Here, we report on LysM effectors of the soil-borne fungal vascular wilt pathogen *Verticillium dahliae*. Comparative genomics revealed three core LysM effectors that are conserved in a collection of *V. dahliae* strains. Remarkably, and in contrast to the previously studied LysM effectors of other plant pathogens, no expression of core LysM effectors was monitored *in planta* in a taxonomically diverse panel of host plants. Moreover, targeted deletion of the individual LysM effector genes in *V. dahliae* strain JR2 did not compromise virulence in infections on Arabidopsis, tomato or *Nicotiana benthamiana*. Interestingly, an additional lineage-specific LysM effector is encoded in the genome of *V. dahliae* strain VdLs17 but not in any other *V. dahliae* strain sequenced to date. Remarkably, this lineage-specific effector is expressed *in planta* and contributes to virulence of *V. dahliae* strain VdLs17 on tomato, but not on Arabidopsis or on *N. benthamiana*. Functional analysis revealed that this LysM effector binds chitin, is able to suppress chitin-induced immune responses, and protects fungal hyphae against hydrolysis by plant hydrolytic enzymes. Thus, in contrast to the core LysM effectors of *V. dahliae*, this lineage-specific LysM effector of strain VdLs17 contributes to virulence *in planta*.

## INTRODUCTION

To establish infection, fungal plant pathogens secrete effector molecules that manipulate host physiology, including immune responses that are triggered when plant hosts sense invading pathogens (Jones and Dangl, 2006; Rovenich et al., 2014; Cook et al., 2015). Typically, effectors are small secreted proteins that are species- or even lineage-specific. However, some effectors are more broadly conserved, such as the necrosis- and ethylene-inducing-like proteins (NLPs) that are produced by bacteria, oomycetes and fungi, and that are particularly known for their phytotoxic activity (de Jonge et al., 2011; Gijzen and Nürnberger, 2006). Another group of conserved fungal effectors are lysin motif (LysM) effectors which are found in a wide range of fungal species, including plant and animal pathogens as well as saprophytes (de Jonge and Thomma, 2009; Kombrink and Thomma, 2013).

LysM effectors are defined as secreted proteins that contain no annotated protein domains apart from a varying number of LysM domains, which are carbohydrate binding modules that occur in many prokaryotic and eukaryotic proteins (de Jonge and Thomma 2009; Buist et al., 2008). The LysM effectors studied to date belong to various fungal plant pathogenic species and were all found to bind chitin, a homopolymer of unbranched β-1-4-linked *N*-acetyl-glucosamine (GlcNAc) (de Jonge et al., 2010; Marshall et al., 2011, Mentlak et al., 2012; Takahara et al., 2016). Chitin is a major component of fungal cell walls and plays an important role in the interaction between fungal pathogens and their plant hosts (Bowman and Free, 2006; Kombrink et al., 2011; Sánchez-Vallet et al., 2015; Rovenich et al., 2016). Plants evolved to recognize chitin as a ‘non-self’ molecule and mount an immune response upon chitin perception in order to stop fungal infection (Felix et al., 1993; Shibuya et al., 1993). Several plasma membrane-localized chitin receptors have been identified in plants that all contain LysM domains (Kaku et al., 2006; Miya et al., 2007; Wan et al., 2008; Cao et al., 2014). LysM effectors of various fungal foliar pathogens, namely the tomato leaf mould fungus *Cladosporium fulvum*, the wheat blotch fungus *Zymoseptoria tritici* (formerly *Mycosphaerella graminicola*), the rice blast fungus *Magnaporthe oryzae* and the Brassicaceae anthracnose fungus *Colletotrichum higginsianum*, were demonstrated to perturb the activation of chitin-induced immunity during host colonization and contribute to virulence (de Jonge et al., 2010; Marshall et al., 2011, Mentlak et al., 2012; Takahara et al., 2016). Recently, the crystal structure of the LysM effector Ecp6, secreted by *C. fulvum*, revealed that Ecp6 suppresses chitin-triggered immunity through two distinct mechanisms (Sánchez-Vallet et al., 2013). One mechanism involves two LysM domains of a single Ecp6 molecule (LysM1 and -3) that cooperatively bind chitin with ultra-high (pM) affinity and allows Ecp6 to outcompete plant receptors for chitin binding. In addition, also the remaining, singular, LysM domain (LysM2) binds chitin and has the capability to suppress chitin-induced immune responses. However, its relatively low (μM) affinity for chitin likely does not allow this domain to function through sequestration of chitin fragments, suggesting that LysM2 suppresses chitin-triggered immunity through another mechanism that potentially involves receptor complex disturbance (Sánchez-Vallet et al., 2013). Additionally, LysM effectors Mg1LysM and Mg3LysM from Z. *tritici* were found to protect fungal hyphae against degradation by plant chitinases (Schlumbaum et al., 1986; Marshall et al., 2011). Although the LysM effectors that are characterized to date are all secreted by foliar pathogens, they are also found in the genomes of soil-borne pathogens that infect host plants through their roots, such as *Fusarium oxysporum* and *Verticillium dahliae* (de Jonge and Thomma, 2009).

*V. dahliae* is a soil-borne fungal pathogen that colonizes the xylem vessels of its host plants, resulting in vascular wilt disease (Fradin and Thomma, 2006; Klosterman et al., 2009; Klimes et al., 2015). *V. dahliae* infects a wide range of dicotyledonous plant species, including economically important crops such as cotton, lettuce and tomato. Genetic resistance has been characterized in tomato, and it was shown that the cell surface localized immune receptor Ve1 confers resistance against strains of *V. dahliae* that belong to race 1 (Fradin et al., 2009). However, Ve1 homologs that may recognize *V. dahliae* are widespread in plants (Zhang et al., 2013; Song et al., 2016). Based on comparative population genomics it was discovered that Ve1 recognizes the race 1-specific effector protein Ave1 (for Avirulence on *Ve1* tomato) in order to activate effector-triggered immunity (de Jonge et al., 2012). Furthermore, *Ave1* is required for full virulence on tomato genotypes that lack *Ve1* (de Jonge et al., 2012). Additionally, comparative population genomics revealed that all *V. dahliae* strains carry lineage-specific genomic regions that account for 1-5 Mb of their ^~^35 Mb genome and that are significantly enriched for *in planta* expressed genes (de Jonge et al., 2012; 2013; Faino et al., 2015; 2016). While in several other plant pathogenic species, such as *F. oxysporum* and *Z. tritici*, such lineage-specific regions are found as small dispensable chromosomes (Ma et al., 2010; Goodwin et al., 2011), in *V. dahliae* these regions are found as islands within the core chromosomes (de Jonge et al., 2013; Faino et al., 2015; 2016).

In this manuscript we characterize the LysM effector gene family of *V. dahliae* that consists of three members in the core genome of *V. dahliae*. Remarkably, one additional LysM effector gene was found in an lineage-specific region of *V. dahliae* strain VdLs17.

## RESULTS

### Three core LysM effectors are encoded in the *V. dahliae* genome

Initially, LysM effectors were identified in the first available *V. dahliae* genome sequence; that of strain VdLs17 (Klosterman et al., 2009). However, as this strain has been isolated from the non-model plant lettuce and is a rather mild pathogen of tomato and Arabidopsis (Yadeta and Thomma, unpublished data), our research focuses on *V. dahliae* strain JR2 that is particularly aggressive on these plant hosts (de Jonge et al., 2012; 2013; Santhanam et al., 2013; 2016; Santhanam and Thomma, 2013). Recently, we established a gapless whole-genome sequence of the *V. dahliae* JR2 strain (Faino et al., 2015; http://fungi.ensembl.org/Verticillium_dahliaejr2/Info/Index). The seven LysM effector genes that were initially identified in strain VdLs17 were also found to occur in JR2, except for the lineage-specific effector VDAG_05180 (Klosterman et al., 2009; de Jonge et al., 2013). Identification of effector genes is often hampered by erroneous gene annotation (Gibriel et al., 2016). Therefore, for the present study we revisisted the corresponding gene models, revealing that not all of the initially identified LysM effector genes were predicted correctly. Firstly, VdLs17 protein VDAG_03096 (VDAG_JR2_Chr8g0580) contains a single LysM domain that only constitutes a minor portion of the total protein (Supplemental Fig. 1). Furthermore, mapping of RNA sequencing reads from samples of *V. dahliae* grown *in vitro* and *in planta* (de Jonge et al., 2012; Faino et al., 2012) indicated that the initially predicted gene model of the corresponding locus is incorrect, as reads mapped to predicted introns while other parts of the predicted gene, including the LysM domain, were not supported by read mapping (Supplemental Fig. 1). Thus, VDAG_03096 (VDAG_JR2_Chr8g05800) was disqualified as a *bona fide* LysM effector. Similarly, deeper analysis of the predicted VdLsl7 VDAG_06426 (VDAG_JR2_Chr7g08210) protein sequence by SMART protein domain analysis (http://smart.embl-heidelberg.de) revealed absence of a signal peptide and presence of a Zinc finger domain (Supplemental Fig. 1), disqualifying this protein as *bona fide* LysM effector. Finally, VdLs17 protein VDAG_08171 (VDAG_JR2_Chr8g01370) was disqualified because of the presence of a glycosylphosphatidylinositol (GPI) anchor, suggesting that it is a membrane-associated protein rather than an apoplastic effector protein. The three remaining VdLs17 LysM effectors VDAG_00902, VDAG_04781 and VDAG_06998 are not only encoded by strain JR2 (as VDAG_JR2_Chrlg02480, VDAG_JR2_Chr6g09650 and VDAG_JR2_Chr4g05910, respectively), but also by 18 additional *V. dahliae* strains that were sequenced (Supplemental Table 1) (de Jonge et al., 2012; 2013; Thomma *et al.*, unpublished data), and are further referred to as core VdLysM effectors. As these effectors comprise four, five or six LysM domains, they were named accordingly; Vd4LysM, Vd5LysM, and Vd6LysM.

**Fig. 1.**
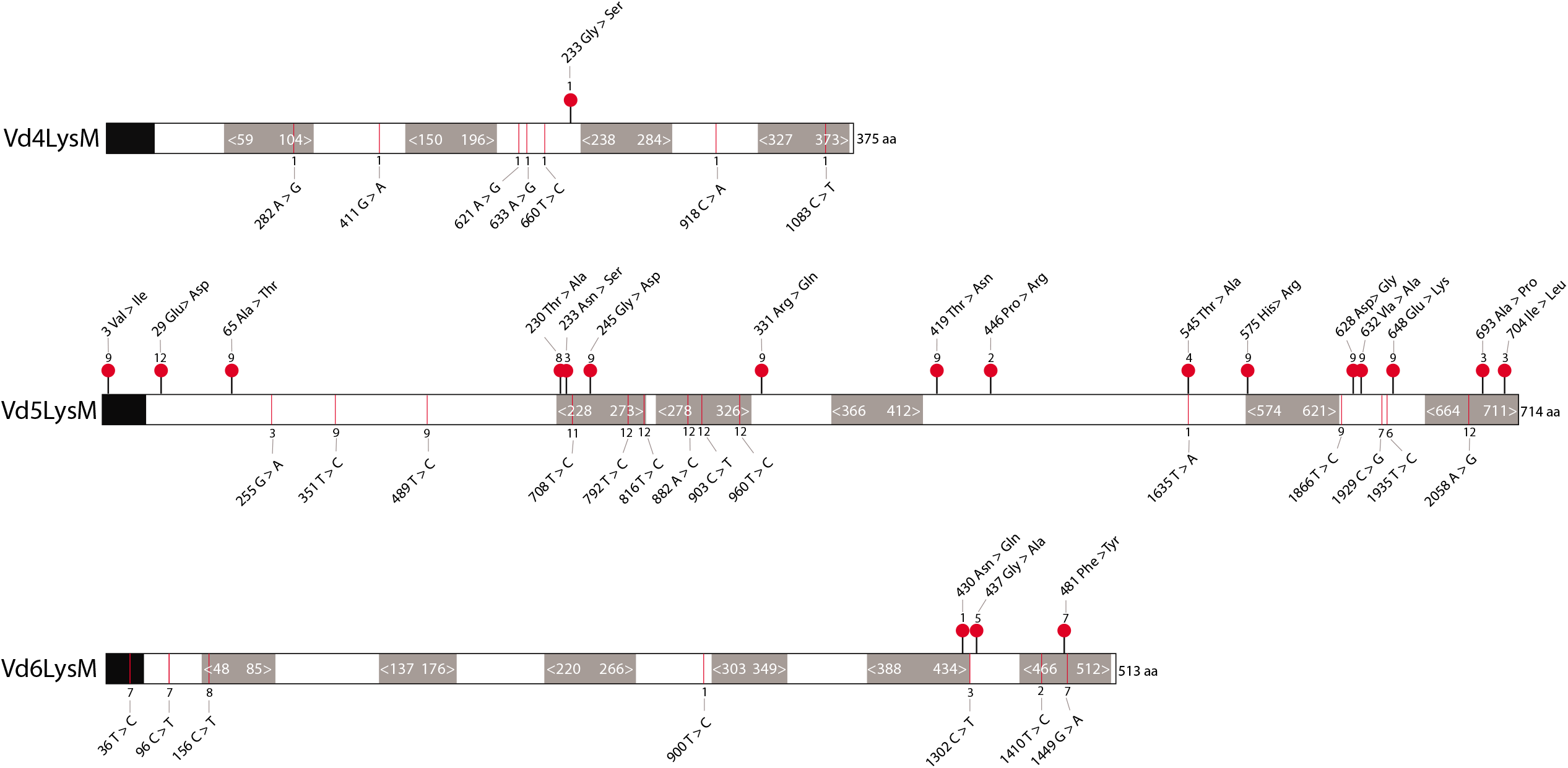
Sequence polymorphisms of core VdLysM effector genes in *Verticillium dahliae* population. The SNPs were compared among 20 *V. dahliae* strains. The protein models are based on the annotation of VdLysMs in *V. dahliae* strain JR2. Three core LysM effectors have four to six predicted LysM domains, which are represented by grey squares (amino acid postions are indicated with white font). The N-terminal black box represents the predicted signal peptide. Sites with synonymous and non-synonymous nucleotide substitutions are marked by red lines and red dots, respectively. The number above or below the SNP sites indicates the amount of strains that share the same nucleotide polymorphism at that position (detailed information on each indivudal SNP can be found in Supplemental Figure 2).

### *VdLysM* sequence analysis in twenty *V. dahliae* strains

To investigate sequence polymorphisms in *VdLysM* effector genes within the *V. dahliae* population, we mapped the sequencing reads of 19 *V. dahliae* strains (Supplemental Table 1) to the genome assembly of strain JR2. Subsequently, we analysed single-nucleotide polymorphisms (SNPs) in the different *V. dahliae* strains. This revealed that *Vd4LysM* is the most conserved LysM effector gene since only one strain (Vd39) showed seven synonymous SNPs and one non-synonymous SNPs (Fig. 1; Supplemental Fig. 2). *Vd5LysM* displays the most SNPs; 14 synonymous and 16 non-synonymous SNPs were found involving the majority of the strains. For *Vd6LysM*, seven synonymous and three non-synonymous SNPs were identified in seven strains in total (Fig. 1; Supplemental Fig. 2). Although the sequence variation in *Vd5LysM* exceeds that of the other *VdLysM* effector genes, no evidence for negative or positive selection could be found. No SNPs were observed in LysM effector genes of strains VdLs17, 2009-605, V152, V52, van Dijk and Vd57 when compared with those of JR2, suggesting that these strains are phylogenetically more closely related to JR2 than the remaining 13 strains.

### Core LysM effectors of *V. dahliae* strain JR2 do not contribute to virulence on tomato

To test whether the core LysM effectors contribute to virulence of *V. dahliae* on tomato, gene functional analysis was pursued in *V. dahliae* strain JR2. Previously, RNA-sequencing was performed on samples harvested during a time course of *V. dahlia*e-inoculated *N. benthamiana* plants (de Jonge et al., 2012; Faino et al., 2012). These data were used to assess the expression of *VdLysM* genes during host invasion. However, no significant *VdLysM* effector gene expression was observed at any of the time points. In contrast, the expression of the functionally analysed *Ave1* effector gene was significantly induced in these samples (de Jonge et al., 2013) (Fig. 2A).

**Fig. 2.**
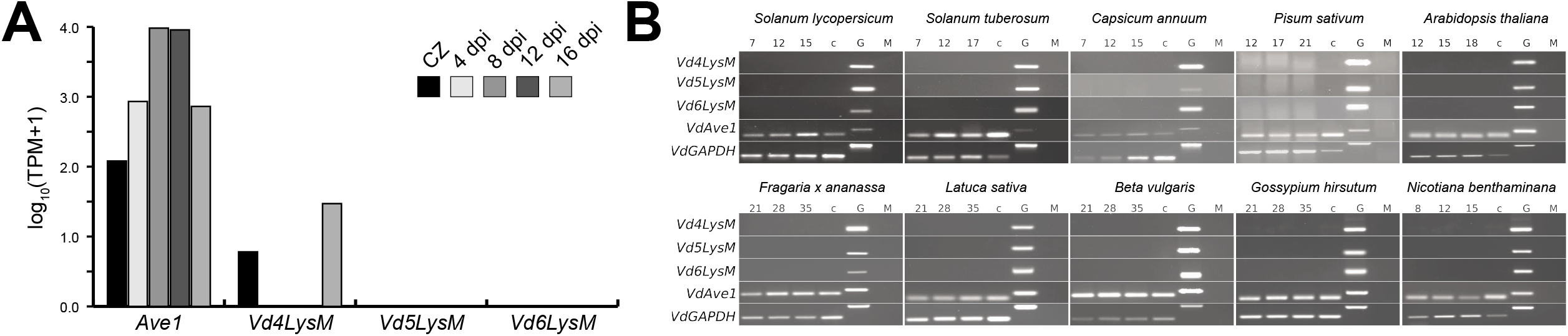
Core VdLysM effector genes of *Verticillium dahliae* strain JR2 are not expressed *in planta.* (A) VdLysM effector gene expression and expression of the *Ave1* effector gene during growth of strain JR2 on Czapek Dox medium (CZ) and during colonization on *Nicotiana benthamiana* at 4, 8,12 and 16 days post inoculation. (B) *In planta* RT-PCR expression analysis of core VdLysM genes. Expression of VdLysM genes could not be detected in ten different plant species (*Solanum lycopersicum, Solanum tuberosum, Capsicum annuum, Pisum sativum, Arabidopsis thaliana, Fragaria* x *ananassa, Latuca sativa, Beta vulgaris, Gossypium hirsutum* and *N. benthaminana*) at timepoints between 7 and 35 dpi. Expression was analyzed at three timepoints per species and referenced to expression of *VdAve1* and *VdGAPDH.* cDNA (c) and genomic DNA (G) from *V. dahliae* cultured *in vitro* were used as controls. Primer pairs were designed to span introns to account for amplification from cDNA or DNA templates. Mock (M) treated plants are also shown.

Differential *V. dahliae* effector gene induction among host plant species has previously been observed, as the *NLP2* effector gene was found to be induced in tomato and Arabidopsis, but not in *N. benthamiana* (Santhanam et al., 2013). Therefore, RT-PCR was performed on cDNA generated from samples of tomato (*Solanum lycopersicum*) and Arabidopsis (*Arabidopsis thaliana*) plants inoculated with *V. dahliae* strain JR2 and harvested at different time points. However, again no expression of any of the *VdLysM* effector genes was recorded, whereas expression of *Ave1* as a positive control was clearly detected (Fig. 2B). In a final attempt to detect *in planta* expression of core *VdLysM* effector genes, a taxonomically wide range of plants that included potato (*Solanum tuberosum*), pepper (*Capsicum annuum*), Australian tobacco (*Nicotiana benthamiana*), pea (*Pisum sativum*), strawberry (*Fragaria* × *ananassa*), lettuce (*Latuca sativa*), beet (*Beta vulgaris*) and cotton (*Gossypium hirsutum*) was inoculated with *V. dahliae* strain JR2 and *VdLysM* effector gene expression was monitored. However, expression of *VdLysM* effector genes could not be detected in any of these plant species, while expression of the *Ave1* effector gene was observed in all hosts (Fig. 2B).

Since it cannot be excluded that *VdLysM* effector gene expression is induced at low levels only, or only at specific time points or at particular sites, further investigations into the possible contribution of the core *VdLysM* effector genes to virulence of *V. dahliae* strain JR2 was evaluated by targeted deletion. Multiple deletion strains were obtained for the LysM effector genes *Vd4LysM, Vd5LysM* and *Vd6LysM*. Importantly, none of the deletion strains that were obtained showed any phenotypic deviations from the wild-type JR2 strain upon cultivation *in vitro* (data not shown). Subsequent virulence assays revealed that Arabidopsis, *N. benthamiana* and tomato plants inoculated with the wild-type strain and those inoculated with deletion strains for *Vd4LysM, Vd5LysM* and *Vd6LysM* showed similar disease development; the timing and degree of stunting of the plants and wilting of the leaves was similar upon inoculation with the various genotypes (Fig. 3). Thus, the lack of *VdLysM* effector gene expression together with the observation that *VdLysM* deletion strains are not affected in virulence on Arabidopsis, *N. benthamiana* and tomato strongly suggest that the core VdLysM effectors do not play an important role during host colonization.

**Fig. 3.**
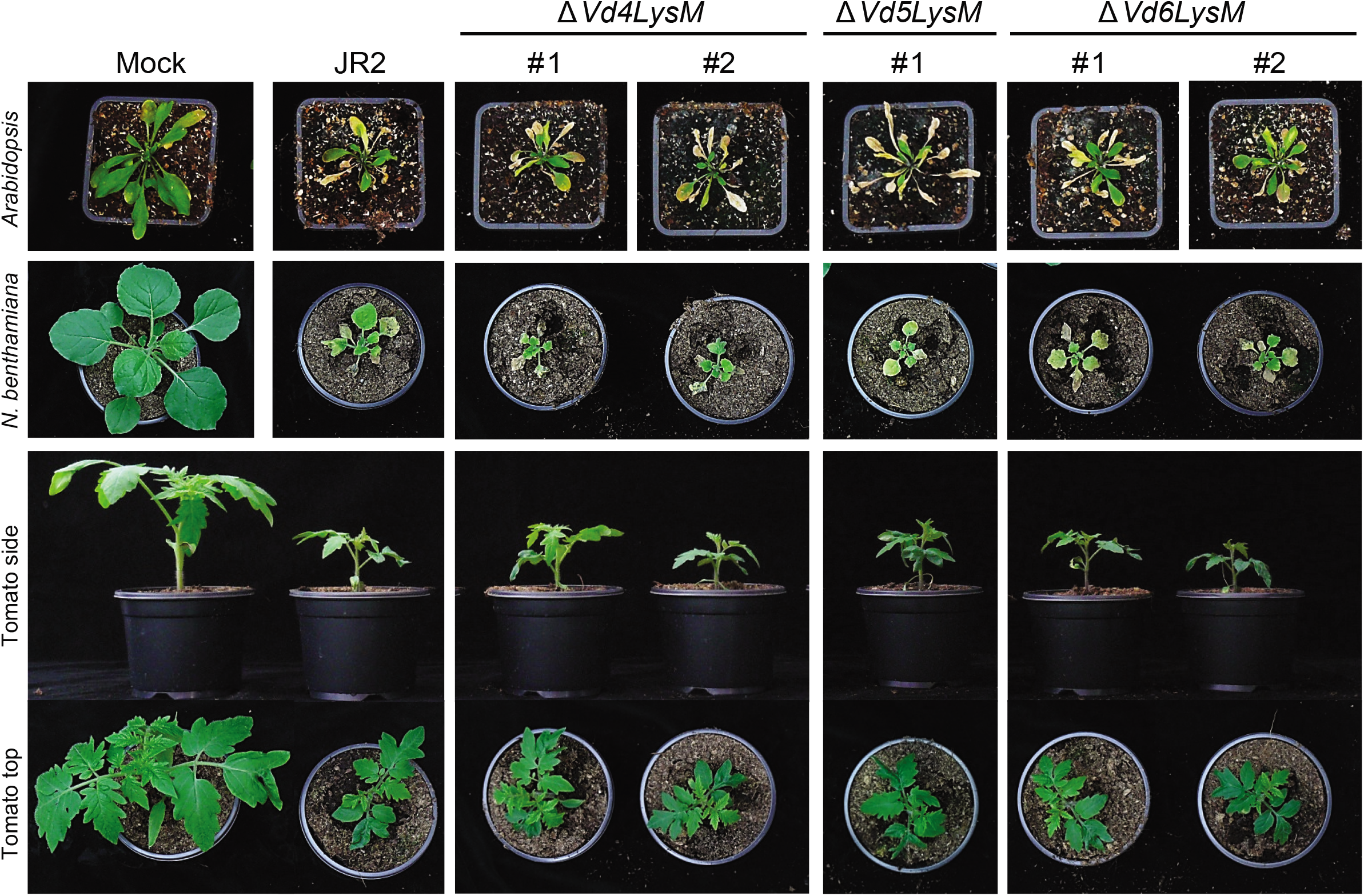
Core VdLysM effector genes do not contribute to virulence of *Verticillium dahliae* strain JR2. Infection assay of wild-type and VdLysM effector gene deletion strains on Arabidopsis, *Nicotiana benthamiana* and tomato plants showing two deletion strains for each VdLysM effector gene, with the exception of *Vd5LysM.* At 14 (tomato, *N. benthamiana*) or 21 (Arabidopsis) days post inoculation pictures were taken of one representative plant out of six that were inoculated with the same *V. dahliae* genotype. The infection assays on Arabidopsis, *N. benthamiana* and tomato were repeated three times with similar results.

### A lineage-specific LysM effector is required for full virulence on tomato, but not on *N. benthamiana* or Arabidopsis

It has previously been demonstrated that lineage-specific effectors of *V. dahliae* strain JR2 contribute significantly to virulence on tomato (de Jonge et al., 2013), suggesting that lineage-specific regions of individual *V. dahliae* genotypes are important for the development of aggressiveness on particular host plants. Intriguingly one LysM effector gene, *VDAG_05180*, was originally identified in strain VdLs17 but could not be found in strain JR2 nor in any of the other 18 additional *V. dahliae* strains that were sequenced, revealing that *VDAG_05180* is a lineage-specific effector gene. Additionally, it was shown that this effector, that contains two LysM domains and is hence designated Vd2LysM, contributes to virulence on tomato (de Jonge et al., 2013). Here, the expression of *Vd2LysM* during infection of *V. dahliae* strain VdLs17 on tomato, *N. benthamiana* and Arabidopsis was investigated using real-time PCR, and expression was detected at each of the time points monitored (Fig. 4A). To test whether Vd2LysM contributes to virulence, *Vd2LysM* deletion strains were inoculated on these host plants. Importantly, none of the transformants showed morphological anomalies upon growth *in vitro* when compared with the VdLs17 wild-type strain (Fig. 4B). As noted previously (de Jonge et al. 2013), the *Vd2LysM* deletion strains showed significantly reduced virulence upon inoculation on tomato when compared with the wild-type strain VdLs17 (Fig. 4C). The tomato plants that were inoculated with the wild-type strain showed stronger stunting than the plants that were inoculated with two independent *ΔVd2LysM* strains (Fig. 4C). In accordance with the reduced symptom development, real-time PCR quantification of fungal biomass showed that the *ΔVd2LysM* strains produced significantly less biomass than the wild-type (Fig. 4E). These results show that the lineage-specific LysM effector gene *Vd2LysM* plays a role in virulence of *V. dahliae*. To investigate whether the contribution to virulence extends to other plant hosts as well, the *Vd2LysM* deletion strains were inoculated onto *N. benthamiana* and Arabidopsis plants. Intriguingly, the virulence of the *Vd2LysM* deletion strains appeared to be comparable to that of the wild-type strain on these plant species, as all genotypes induced similar symptomatology on the inoculated plants and real-time PCR measurements of fungal biomass did not reveal compromised fungal host colonization (Fig. 4D & Fig 4E). On Arabidopsis, it even seems that *Vd2LysM* deletion leads to enhanced colonization.

**Fig. 4.**
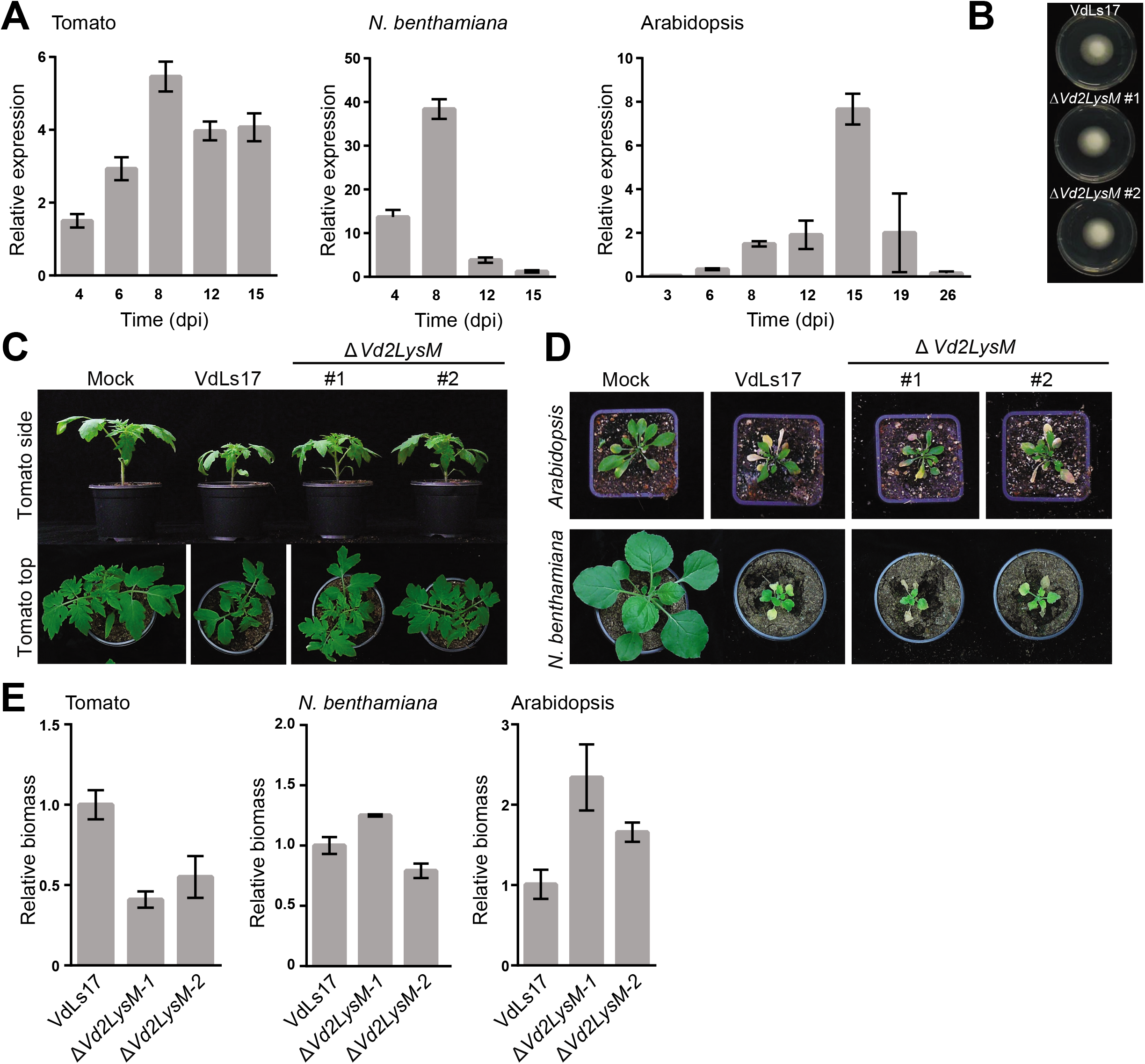
Lineage-specific LysM effector Vd2LysM contributes to virulence of *Verticillium dahliae* strain VdLsl7 on tomato. (A) *Vd2LysM* expression during colonization of *V. dahliae* strain VdLsl7 on tomato, *N. benthamiana* and Arabidopsis plants at 2-26 days post inoculation (DPI). (B) Morphology of wild-type *V. dahliae* strain VdLsl7 and two *Vd2LysM* deletion strains after 7 days of incubation on PDA medium at room temperature. (C) Pictures of representative tomato plants out of eight plants that were either mock-inoculated or inoculated with wild-type *V. dahliae* strain VdLs17 and two deletion strains of *Vd2LysM* at 14 DPI. The infection assay was repeated tree times with similar results. (D) Pictures of representative *N. benthamiana* and Arabidopsis plants out of eight plants that were either mock-inoculated or inoculated with wild-type *V. dahliae* strain VdLsl7 and two deletion strains of *Vd2LysM* at 14 DPI (*N. benthamiana*) and 21 (Arabidopsis) DPI. The assay was repeated tree times with similar results. (E) Fungal biomass accumulation in tomato, *N. benthamiana* and Arabidopsis plants inoculated with wild-type strain VdLs17 and Vd2LysM deletion strains at 14 (tomato, *N. benthamiana*) and 21 (Arabidopsis) DPI. Error bars represent the standard error of three replicate experiments.

### Vd2LysM binds chitin

Previously characterized LysM effectors of fungal plant pathogens were demonstrated to bind chitin (de Jonge et al., 2010; Marshall et al., 2011; Mentlak et al., 2012; Takahara et al., 2016). Therefore, the chitin-binding ability of Vd2LysM was tested. To this end, Vd2LysM was heterologously produced in the yeast *Pichia pastoris*, which has previously been used for production of LysM effectors from other fungi (Kombrink, 2012). Subsequently, the purified Vd2LysM protein was used in affinity precipitation assays with the insoluble carbohydrates chitin, chitosan, xylan and cellulose. We observed that Vd2LysM precipitated with all carbohydrates tested (Fig. 5A), which suggests that the protein precipitates by itself rather than binds to any of these carbohydrates. To investigate this further, Vd2LysM was subjected to glycan-array analysis to test binding affinity for ^~^600 glycans, taking *C. fulvum* Ecp6 as a control. As demonstrated previously, Ecp6 specifically bound to the chitin oligosaccharides (GlcNAc)_3_, (GlcNAc)_5_ and (GlcNAc)_6_ that are present on the array (de Jonge et al., 2010). In contrast, Vd2LysM did not bind to any of the glycans on the array (Supplemental Fig. 3). Based on these findings we concluded that *P. pastoris*-produced Vd2LysM likely precipitated spontaneously in the affinity precipitation assay.

**Fig. 5.**
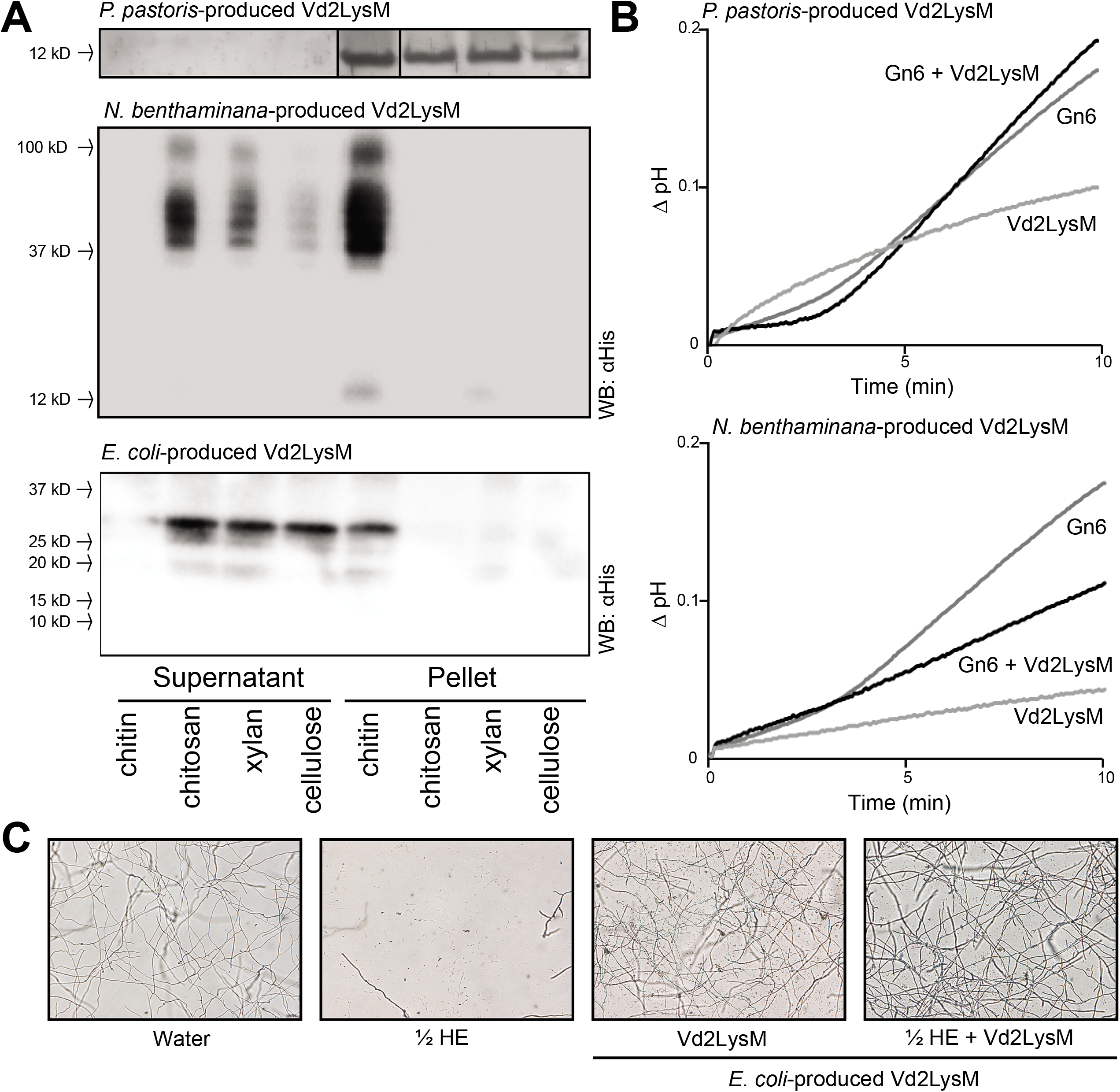
Vd2LysM is a chitin-binding protein that suppresses chitin-induced immune responses. (A) Vd2LysM produced in *Pichia pastoris, Nicotiana benthamiana* and *Escherichia coli* was used in affinity precipitation experiments with insoluble polysaccharides chitin, chitosan, xylan or cellulose. After SDS-PAGE and Coomassie Brilliant blue staining for *P. postoris*-produced Vd2LysM or western blot analysis for *in* planta-and *E. coli*-produced Vd2LysM, the proteins are observed in the insoluble pellet fraction or in the supernatant fraction. Vd2LysM produced in *P. pastoris* is observed in the insoluble pellet fraction of all polysaccharides. Vd2LysM produced *in planta* and in *E coli* precipitates with chitin but not with chitosan, xylan or cellulose. (B) pH-shift measurements in a tomato cell suspension after addition of 1 nM chitin or 1 nM chitin that is pre-incubated with 10 nM Vd2LysM produced in either *P. pastoris* or *in planta*. The graphs show a representative experiment out of two to four experiments with similar results. (C) Micrographs of *Fusarium oxysporum* f.sp. *lycopersicum* strain 4287 taken 23 hours after the addition of water, hydrolytic enzyme mix (HE), or Vd2LysM and pre-treated with Vd2LysM followed by treatment of hydrolytic enzyme mix. The figure is representative of three independent experiments.

Recently, a crystal structure was obtained of *C. fulvum* Ecp6 that was heterologously produced in *P. pastoris*. Unanticipated, the crystal structure of the protein contained chitin, arguably derived from *P. pastoris* during protein production, in a composite binding groove of two LysM domains (Sánchez-Vallet et al., 2013). To exclude the possibility that potential Vd2LysM substrate binding sites were occupied by *P. pastoris* chitin, the effector was produced in *N. benthamiana*. The purified protein was subsequently tested for chitin-binding ability in an affinity precipitation assay using chitin, chitosan, xylan and cellulose. Interestingly, *N. benthamiana*-produced Vd2LysM bound to chitin, but not to any of the other carbohydrates (Fig. 5A). Furthermore, we observed that *N. benthamiana*-produced Vd2LysM migrates as multiple bands on gel, which were not visible with *P. pastoris*-produced Vd2LysM that migrates as a single band of the expected 17 kDa. This may indicate that Vd2LysM produced in *N. benthamiana* forms oligomers. Collectively, these findings suggest that Vd2LysM is a chitin-binding effector protein, and the chitin that is readily available during heterologous production in *P. pastoris* occupied the Vd2LysM substrate binding sites.

### Vd2LysM suppresses chitin-induced immune responses and protects hyphae against degradation by plant hydrolytic enzymes

Previously, LysM effectors from various fungal plant pathogens were demonstrated to suppress the chitin-induced pH-shift in a tomato cell suspension, which is indicative for the ability of the effector to perturb chitin-induced host immune responses (Felix et al., 1993; de Jonge et al., 2010, Marshall et al., 2011, Mentlak et al., 2012; Takahara et al., 2016). Both *in planta*-produced and *P. pastoris*-produced Vd2LysM were tested for this capacity as described previously (de Jonge et al., 2010). Interestingly, *P. pastoris*-produced Vd2LysM that is not able to bind chitin did not suppress the chitin-induced pH-shift, whereas *N. benthamiana* produced Vd2LysM that is able to bind chitin, suppressed the chitin-induced immune response (Fig. 5B). These results suggest that Vd2LysM is able to suppress chitin-triggered immune responses during *V. dahliae* colonization of tomato.

As the Z. *tritici* LysM effectors Mg3LysM and Mg1LysM were previously demonstrated to protect fungal hyphae against degradation by plant hydrolytic enzymes (Marshall et al., 2011), we also intended to test Vd2LysM for this activity. However, the yield of *in planta*-produced Vd2LysM was too low to perform such assays. Therefore, we pursued Vd2LysM production in yet another heterologous system; *Escherichia coli*. Similar to *N. benthamiana*-produced Vd2LysM, also *E. coli*-produced Vd2LysM was found to bind chitin but no other insoluble carbohydrates tested (Fig. 5A). Moreover, also *E. coli*-produced Vd2LysM showed signs of oligomerization. Interestingly, similar to the Z. *tritici* LysM effectors Mg3LysM and Mg1LysM, also Vd2LysM was found to protect fungal hyphae against degradation by plant hydrolytic enzymes (Fig. 5C).

### LysM domains of Vd2LysM are more similar to LysM domains of previously characterized LysM effectors than to those of core VdLysM effectors

Vd2LysM is presently the only LysM effector of *V. dahliae* for which a role in virulence could be shown, and that is able to perturb chitin-centred host immune responses as previously described for Ecp6, Mg1LysM, Mg3LysM, Slp1, ChELP1 and ChELP2 (de Jonge et al., 2010; Marshall et al., 2011; Mentlak et al., 2012; Takahara et al., 2016). Remarkably, the LysM domains of Vd2LysM are more similar to LysM domains of Ecp6, Mg1LysM, Mg3LysM, Slp1, ChELP1 and ChELP2 than to the LysM domains of the core VdLysM effectors (Fig. 6). To further investigate this, we carried out a comparative LysM domain analysis with functionally characterized plant and fungal proteins (Fig. 6). The fungal LysM domains fall into two separate clades (Fig. 6B). One clade was nearly exclusively formed by LysMs of plant receptors and LysMs of fungal effectors with a role in virulence, including Vd2LysM (Bolton *et al.*, 2008; Marshall *et al.*, 2011; Mentlak *et al.*, 2012; Takahara et al., 2016). This suggests that fungi and plants produce LysM proteins with a conserved motif and, thus, possibly originate from a common ancestor. Interestingly, the second clade solely contains the LysMs of the *V. dahliae* core effectors and the *Trichoderma atroviride* LysM effector TAL6 that was previously shown to specifically inhibit germination of conidia of *Trichoderma* spp. and that was proposed to act in fungal development rather than in host interactions (Seidl-Seiboth et al., 2013) (Fig. 6B). Intriguingly, the two clades are characterized by highly divergent consensus motifs for their LysMs, with the ‘fungal-specific’ LysMs having three highly conserved cysteine residues that are completely lacking from the ‘effector/receptor’ LysMs. Collectively, these findings may suggest that the core LysM effectors of *V. dahliae*, in contrast to Vd2LysM, do not act in host interactions, but possibly in other physiological processes.

**Fig. 6.**
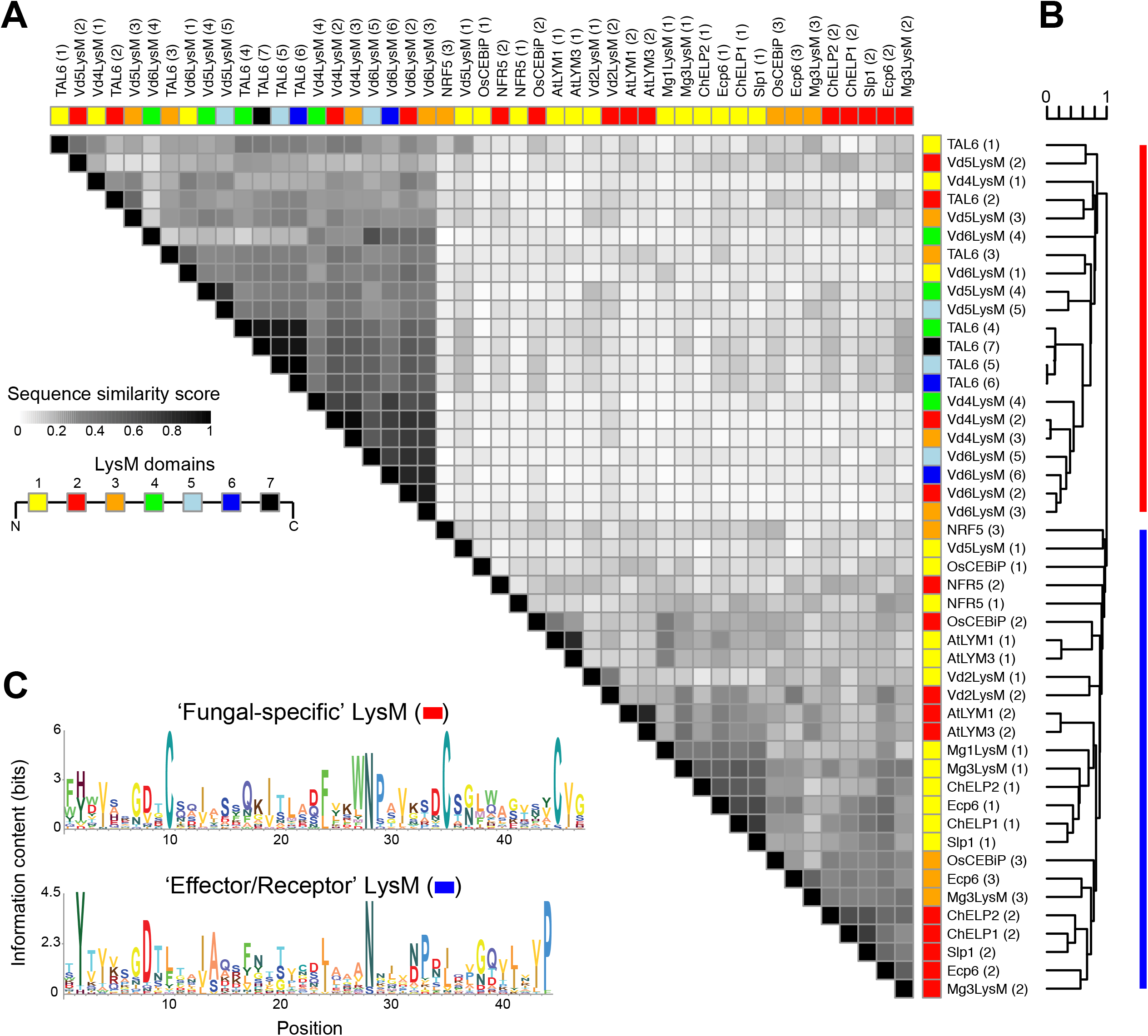
Sequence similarity between LysM domains. (A) Pairwise comparisons of LysM domains of fungal and plant LysM proteins. Normalized sequence similarities between LysM domains are displayed in the heat map. Colour annotation above the heat map indicates the relative position of each LysM domain within the LysM protein. (B) Hierarchical clustering (UPGMA clustering) based on the (dis)-similarity matrix of all pairwise LysM domain comparisons are shown above and on the side, and the clustering is used to order the rows and columns of the heat map. (C) Sequence logos of LysM domains identified in fungal effectors as well as in plant receptors (blue) and LysM domains that have been previously defined as ‘fungal-specific’.

## DISCUSSION

Here we describe the characterization of the LysM effector catalogue of the soil-borne vascular wilt pathogen *V. dahliae*. In-depth analysis revealed that three *VdLysM* effector genes occur in the core genome of all strains for which a genome sequence is available. In addition, a lineage-specific *VdLysM* effector gene, *Vd2LysM*, occurs in strain VdLs17 and not in any of the other strains that have been sequenced to date. We initially reasoned that the ubiquitous occurrence of the three core LysM effector genes suggests that these are required for host colonization by *V. dahliae.* However, a role in virulence cannot be attributed to any of these core LysM effectors. Intriguingly, the lineage-specific Vd2LysM effector was found to contribute to host-specific virulence of strain VdLs17.

LysM effectors are found in fungal species with various lifestyles, but thus far have only been characterized as virulence factors of plant pathogenic fungi (de Jonge et al., 2010; Marshall et al., 2011; Mentlak et al., 2012; Takahara et al., 2016). Moreover, they were only studied in foliar pathogens with a (relatively) narrow host range: *C. fulvum* (Ecp6), *Z. tritici* (Mg1LysM and Mg3LysM), *M. oryzae* (Slp1) and *C. higginsianum* (ChELP1 and ChELP2) (de Jonge et al., 2010; Marshall et al., 2011; Mentlak et al., 2012; Takahara et al., 2016). In contrast, *V. dahliae* is a soil-borne vascular plant pathogen that infects a broad range of host plants. The expression of the previously characterized LysM effectors is induced during the early stages of infection of the respective pathogens, when evasion of recognition by the host is in particular important to facilitate tissue colonization. The expression of *Vd2LysM* peaks at about one week after inoculation, which is when *V. dahliae* biomass is accumulating in the xylem and before wilting and necrosis of host tissue is visible (Fig. 4A) (Fradin and Thomma, 2006). Ecp6, Mg3LysM, Slp1, ChELP1 and ChELP2 were demonstrated to function as suppressors of chitin-induced host immunity and, similar to these proteins, Vd2LysM suppresses chitin-induced medium alkalinisation in a tomato cell suspension. This strongly suggests that Vd2LysM contributes to virulence of *V. dahliae* through perturbing the activation of chitin-triggered host immunity. However, in contrast to Ecp6, Slp1, ChELP1 and ChELP2, and similar to Mg3LysM, Vd2LysM is also able to protect fungal hyphae against degradation by plant hydrolytic enzymes. Thus, Vd2LysM plays a broad role in overcoming chitin-centred host immune responses (Sánchez-Vallet et al., 2015).

Considering the widespread occurrence of chitin receptor homologues in plant species, and the observation that LysM effectors of diverse plant pathogens have the ability to suppress chitin-triggered immunity, it seems that this ability is fundamental for fungal plant pathogens to establish infections on their hosts. As most of the sequenced *V. dahliae* strains are pathogenic on tomato, these strains likely employ other molecules than Vd2LysM to overcome chitin-centred host immune responses (Sánchez-Vallet et al., 2015). Alternatively, *V. dahliae* may employ other mechanisms to prevent the activation of chitin-triggered immune responses (Sánchez-Vallet et al., 2015; Rovenich et al., 2016). For example, the α-1,3-glucan synthase gene was demonstrated to be important for virulence of several fungal pathogens, as the α-1,3-glucan layer around hyphae reduces the accessibility of fungal chitin to host chitinases, and consequently prevents release of free chitin fragments (Fujikawa et al., 2012). In addition, fungal species were found to secrete chitin deacetylases that convert chitin into chitosan, which is a poor inducer of immune responses (Gough and Cullimore, 2011).

Structural analysis of the *C. fulvum* LysM effector Ecp6 revealed that the first and third LysM domain cooperatively bind one chitin molecule with ultra-high (pM) affinity through intramolecular LysM dimerization, whereas the second, singular LysM domain displays low micromolar affinity (Sánchez-Vallet et al., 2013). Nevertheless, both binding sites are able to suppress chitin-triggered immune responses (Sánchez-Vallet et al., 2013). Intriguingly, all characterized LysM effectors that have the ability to suppress chitin-triggered immune responses either have three (Ecp6, Mg3LysM) or two (Slp1, ChELP1, ChELP2, Vd2LysM) LysM domains. It remains presently unknown whether the LysM effectors with two LysM domains contain one composite chitin binding site that is composed by the coordinated action of two LysM domains, or two binding sites of separate, singular LysM domain activities. Consequently, the mechanism by which Vd2LysM perturbs the activation of chitin-triggered immunity needs further investigation.

Recently, evolutionary analyses and consensus pattern profiling of fungal LysM domains led to their classification into two groups (Akcapinar et al., 2015). The clustering into two phylogenetic clades across taxonomic boundaries was consistent with the presence of a characteristic cysteine residue pattern present in 80% of fungal-specific LysMs (Akcapinar et al., 2015) (Fig 6C). Remarkably, the group of fungal LysMs, which do not display the conserved cysteine residues, contains all of the LysM domains of previously characterized LysM effectors from fungal pathogens (Bolton et al., 2008; Marshall et al., 2011; Mentlak et al., 2012; Takahara et al., 2016) (Fig 6C). It was previously noted that LysM domains from saprotrophs, mutualists and mycoparasitic fungi were underrepresented in this clade (Akcapinar et al., 2015), pointing toward significantly diverging roles of LysM proteins. To predict the putative roles of the core LysM effectors of *Verticillium*, we carried out a comparative LysM domain analysis with functionally characterized plant and fungal proteins (Fig. 6). Consistent with previous findings (Akcapinar *et al.*, 2015), the fungal LysM domains analysed here largely fall into two separate clades (Fig. 6B). Interestingly, the *V. dahliae* core effectors and TAL6 from the mycoparasitic fungus *T. atroviride*, which selectively inhibits spore germination of *Trichoderma* spp. (Seidl-Seiboth et al., 2013), grouped into same clade (Akcapinar *et al.*, 2015) (Fig 6B). Considering the similarity of the core LysM effectors of *V. dahliae* to TAL6, we hypothesize that they could function in other physiological processes rather than virulence. Although the (single gene) mutants do not display phenotypic deviations from the wild type strain, further experimental support is required to confirm this hypothesis. Originally, the LysM domain was identified in bacterial lysozymes that bind and hydrolyse peptidoglycan components of bacterial cell walls (Buist et al., 2008). Therefore, it is possible that also fungal LysMs display peptidoglycan-specific binding activities, which could help fungi to outcompete bacterial competitors (Kombrink & Thomma, 2013). Similar to the LysM effector functions of plant-colonizing fungi, LysM effectors of saprotrophs could protect fungal hyphae from hydrolytic enzyme attack secreted by mycoparasites.

In conclusion, we demonstrate that a lineage-specific LysM effector of *V. dahliae* contributes to virulence, whereas three LysM effectors that are present in the core genome do not seem to play a role during host colonization. This finding confirms previous observations that especially those effectors that are encoded in lineage-specific regions of *V. dahliae* genomes are important for fungal aggressiveness (de Jonge et al., 2013). It is tempting to speculate that the core LysM effectors play other roles in fungal life. Therefore, it might be worthwhile to investigate whether the core LysM effectors contribute to fungal growth in other stages of the *V. dahliae* life cycle that were not covered in this study, for example during saprophytic growth or during survival as resting structure in the soil.

## EXPERIMENTAL PROCEDURES

### VdLysM effector gene expression analysis

Conidiospores of *V. dahliae* strain JR2 were inoculated on potato dextrose agar plates and grown at room temperature for 7 days before harvest and dilution in 40 ml of potato dextrose broth to a concentration of 1×10^6^ conidia/ml that was used as inoculum for plant inoculation. Plants from S. *lycopersicum, S. tuberosum, C. annuum, P. sativum, A. thaliana, F. x ananassa, L. sativa, B. vulgaris, G. hirsutum* and *N. benthaminana* were grown at the greenhouse at 21°C/19°C during 16 h/8 h light/dark photoperiods, respectively, with a relative humidity of ^~^75%, and 100 W/m^2^ supplemental light when light intensity dropped below 150 W/m^2^. At 10 days post germination, the plants were inoculated with *V. dahliae* or PDB as mock treatment by dipping of the roots of uprooted plants for 6 min before transfer to fresh soil. Three time points were set for each species and five inoculated plants were collected per time point.

A two-step RT-PCR was performed to test the expression of VdLysM effectors. RNA was extracted from all sampled plant material using the QuickRNA™ MiniPrep kit (Zymo Research, Irvine, CA, USA) following the protocol provided by the manufacturer. cDNA was synthetized from 1 μg of RNA from each sample by M-MMLV reverse transcriptase (Promega, Madison, WI, USA) according to the manufacturer’s instructions. cDNA and genomic DNA from *V. dahliae* strain JR2 grown *in vitro* on PDB were used as controls. Expression analysis was performed by PCR using 2 μl of cDNA as template with the following protocol: 5 min at 95°C followed by 32 cycles of 95°C for 30 seconds, 58°C for 30 seconds and 72°C for one minute. For the strawberry, lettuce, beet and cotton samples, a second PCR was needed to detect expression of the control genes, owing to the low levels of *V. dahliae* biomass in infected plants. This second PCR was performed by taking 2 μl of the first PCR reaction as template and run with the same temperature profile, but for only 15 cycles instead. Primers listed in Supplemental Table 2 were used to amplify transcripts from *Vd4LysM, Vd5LysM* and *Vd6LysM. VdAVe1* and *VdGAPDH* were used as controls for effector induction *in planta* and presence of *V. dahliae* in the sample, respectively. Primers were designed to span introns to discriminate between amplicons derived from cDNA or DNA templates as a control for DNA contaminations in the synthesized cDNA. Products from all samples were visualized by gel electrophoresis in a 2% agarose gel.

### Functional analysis of *VdLysM* effector genes

*VdLysM* effector gene deletion strains were generated by amplifying flanking sequences of the coding sequences using the primer sets listed in Supplemental Table 2. PCR products were subsequently cloned into pRF-HU2 (Frandsen et al. 2008). *V. dahliae* transformation and subsequent inoculations on tomato (cv. Motelle and MoneyMaker) plants to assess the virulence of the knockout mutants were performed as described (Fradin et al. 2009). In one experiment, six to eight plants were used per inoculation with wild-type or deletion strains and the experiment was repeated at least three times. Plants were regularly inspected and representative plants were photographed at 12 and 21 days post-inoculation (DPI). For biomass quantification, the roots and stem below the cotyledons of four plants per *V. dahliae* genotype were flash-frozen in liquid nitrogen. The samples were ground to powder, of which an aliquot was used for DNA isolation (Fulton et al. 1995). Real-time PCR was conducted with primer sets SlRub-F1/SlRub-F2 for tomato *RuBisCo* and VdGAPDH-F/VdGAPDH-R for *V. dahliae GAPDH* (Supplemental Table 2). For expression analyses, 3-week-old *N. benthamiana* plants were inoculated with strain VdLs17 as previously described (Fradin et al. 2009), harvested at 4, 6, 8 and 12 DPI, and flash-frozen in liquid nitrogen. Total RNA was extracted using the RNeasy Kit (Qiagen, Hilden, Germany) and cDNA was synthesized by SuperScript III (Invitrogen, Carlsbad, CA, USA). Real-time PCR was conducted with primer sets Q-VdGAPDH-F/Q-VdGAPDH-R for *V. dahliae GAPDH* and Q-Vd2LysM- F/Q-Vd2LysM-R for *V dahliae Vd2LysM* (Supplemental Table 2).

### *VdLysM* sequence analysis

Sequencing reads of 19 *V. dahliae* stains (Supplemental Table 1) were mapped to the genome assembly of strain JR2 using BWA with default settings (Li and Durbin, 2009). Subsequently, SNPs in genes encoding core LysM proteins were detected using FreeBayes with default settings (Garrison and Marth, 2012). The effects of SNPs were predicted using VEP (McLaren et al., 2010).

### Heterologous Vd2LysM expression in *Pichia pastoris*

*Vd2LysM* was cloned into *P. pastoris* expression vector pPic9 (Invitrogen, Carlsbad, CA, USA) after performing PCR using primers to add the N-terminal HIS- and FLAG-tag and *Eco*RI and *Not*I restriction sites for directional cloning (Supplementary Table 1). Subsequently, *P. pastoris* strain GS115 was transformed and a selected clone was cultured in a fermentor (Bioflo 3000) as described previously (Rooney et al., 2005). After removal of cells and concentration of the culture medium the HIS-tagged protein was purified using a Ni-NTA column (Qiagen, Hilden, Germany) according to the manufacturer’s protocol. The final protein concentration was determined spectrophotometrically at 280 nm and confirmed using the Pierce BCA Protein Assay Kit (Thermo Scientific, Waltham, MA, USA) with bovine serum albumin (BSA) as a standard.

### *In planta* production of Vd2LysM

PCR was performed to add *Nhe*I and *Sac*I restriction sites to *Vd2LysM*. Using directional cloning, *Vd2LysM* was cloned into vector pHYG, a modified version of the expression vector pMDC32 (Curtis and Grossniklaus, 2003). The expression vector was subsequently transferred into *Agrobacterium tumefaciens* strain MOG101. Infiltration of four- to five-week-old *Nicotiana benthamiana* with *Agrobacterium tumefaciens* was performed as described (Westerhof et al., 2012). Four days after infiltration, leaves were harvested and flash-frozen in liquid nitrogen. Plant material was ground and homogenized in ice-cold extraction buffer (1% v/v Tween-20, 2% w/v immobilized polyvinylpolypyrrolidone (PVPP), 300 mM NaCl, 50 mM NaH_2_PO_4_, 10 mM imidazole, 1 mM Dithiothreitol (DTT), pH = 7,4) using 2 ml/g fresh plant material. After 30 min of homogenizing at 4°C, the crude extract was centrifuged at 16.000 × *g* at 4°C. The supernatant was further cleaned using a miracloth filter, and a Ni-NTA Superflow column (Qiagen, Hilden, Germany) was used to purify the HIS tagged protein, according to the manufacturer’s protocol. The final protein concentration was determined spectrophotometrically at 280 nm and confirmed using the Pierce BCA Protein Assay Kit (Thermo Scientific, Waltham, MA, USA) with bovine serum albumin (BSA) as a standard.

### E. *coli* production of Vd2LysM

The coding sequence of Vd2LysM was amplified from *V. dahliae* cDNA and cloned into the pETSUMO (Invitrogen, Carlsbad, CA, USA) expression vector according to manufacturer’s instructions prior to *E. coli* Origami (DE3) transformation. A single transformant was selected and grown in Luria Broth medium until OD600=0.9. Heterologous production of Vd2LysM was induced with 1 mM IPTG at 26°C during ^~^20 hours. Cell pellets were lysed using lysozyme from chicken egg (Sigma, Saint Louis, MO, USA) and Vd2LysM was purified from the soluble protein fraction using a Ni2+−NTA Superflow column (Qiagen, Hilden, Germany). Purified protein was dialysed against 200 mM NaCl and concentrated over Amicon ultracentrifugal filter units (MWCO=3 kDa; Millipore).

### Sequence similarity between LysM domains

LysM domains within fungal and plant proteins were identified using SMART database (including PFAM domain and Outlier detection) (Schultz et al., 1998). Sequence similarities between LysM domains was approximated using the Needleman-Wunsch algorithm, which has been implemented in the EMBOSS package (Rice et al., 2000). The Needleman-Wunsch score was further normalized to the range 0 to 1 by NW_ij_norm = NW_ij_/max(NW_ii_,NW_jj_). Normalized Needleman-Wunsch scores were displayed in R. Moreover, hierarchical clustering (UPGMA) based on the (dis)-similarity matrix of all pairwise LysM domain was performed in R. Sequence logos for effector/receptor LysM domains and other fungal LysM domain were created with Skylign (Wheeler et al., 2014). LysM domain sequences were aligned using mafft (global setting) (Katoh et al., 2002).

### Affinity precipitation assays

These assays were performed as described (van den Burg et al., 2006) with 100 μg/ml Vd2LysM. The protein was incubated at RT for 1 hr with 3 mg of insoluble polysaccharides while gently rocking.

## ACKNOWLEDGEMENTS

MFS and BPHJT are supported by a Veni and a Vici grant, respectively, of the Research Council for Earth and Life sciences (ALW) of the Netherlands Organization for Scientific Research (NWO). The authors declare no conflict of interest.

## FIGURE LEGENDS

Supplemental Fig. 1. Disqualification of two previously identified *Verticillium dahliae* LysM effector genes. Two originally identified VdLysM effector genes are not predicted correctly. The initially predicted gene model of LysM effector gene *VDAG_03096* is not supported by mapping of RNA sequencing reads (in red). Reads map to predicted introns (in yellow), whereas some coding parts of the gene (in green), including the LysM domain that constitutes only a small part of the predicted protein, is not supported by reads. SMART prediction using the amino acid sequence encoded by VDAG_06426 reveals absence of a signal peptide and presence of a Zinc finger domain. Also in this case, the LysM domain constitutes only a small portion of the predicted protein.

Supplemental Fig. 2. Sequence polymorphisms of core VdLysM effectors in the *V. dahliae* population. Synonumous and non-synonymous polymorphisms and their occurrence in 19 *V. dahliae* strains are indicated per VdLysM effector gene, and nucleotide and amino acid postions of polymorphisms are indicated, respectively.

Supplemental Fig. 3. Glycan array analysis of *Pichia pastoris*-produced Vd2LysM and Ecp6. Relative fluorescence upon scanning of a glycan array that contains probes for 406 glycans after hybridization with Vd2LysM and with Ecp6 as control. Only Ecp6 hybridizes to the array, and only to 170 to 172 representing (GlcNAc)_5_, (GlcNAc)_5_, and (GlcNAc)_3_, respectively.

